# An AI-Driven Framework for Discovery of BACE1 Inhibitors for Alzheimer’s Disease

**DOI:** 10.1101/2024.05.15.594361

**Authors:** Evan Xie, Karin Hasegawa, Georgios Kementzidis, Evangelos Papadopoulos, Bertal Huseyin Aktas, Yuefan Deng

**Author notes:** E-mail: huseyin.

## Abstract

Alzheimer’s Disease (AD) is a progressive neurodegenerative disorder that affects over 51 million individuals globally. The *β*-secretase (BACE1) enzyme is responsible for the production of amyloid beta (A*β*) plaques in the brain. The accumulation of A*β* plaques leads to neuronal death and the impairment of cognitive abilities, both of which are fundamental symptoms of AD. Thus, BACE1 has emerged as a promising therapeutic target for AD. Previous BACE1 inhibitors have faced various issues related to molecular size and blood-brain barrier permeability, preventing any of them from maturing into FDA-approved AD drugs. In this work, a generative AI framework is developed as the first AI application to the *de novo* generation of BACE1 inhibitors. Through a simple, robust, and accurate molecular representation, a Wasserstein Generative Adversarial Network with Gradient Penalty (WGAN-GP), and a Genetic Algorithm (GA), the framework generates and optimizes over 1,000,000 candidate inhibitors that improve upon the bioactive and pharmacological properties of current BACE1 inhibitors. Then, the molecular docking simulation models the candidate inhibitors and identifies 14 candidate drugs that exhibit stronger binding interactions to the BACE1 active site than previous candidate BACE1 drugs from clinical trials. Overall, the framework successfully discovers BACE1 inhibitors and candidate AD drugs, accelerating the developmental process for a novel AD treatment.

## Introduction

Alzheimer’s Disease (AD) is a progressive neurodegenerative disorder that affects over 51 million individuals globally.^1^ In 2019, it was reported as the seventh leading global cause of death.^2^ With the aging human population, these figures are expected to rise before 2060. ^3^ Hence, it is of utmost importance to develop novel drugs to prevent and treat AD.

While the pathology behind AD onset is not fully understood, the predominant hypothesis states that amyloid beta (A*β*) plaques accumulate in the brain, triggering a cascade of events that result in neuronal death and impaired cognitive abilities. ^4^ These A*β* plaques are formed by A*β* peptides, fragments of the healthy Amyloid Precursor Protein (APP) that are produced by *β*-secretase (BACE1) and *γ*-secretase cleavage. Since BACE1 cleavage is the first step in the amyloidogenic processing of APP, it is the rate-limiting step of A*β* production.^5^ Thus, the BACE1 protein has become a promising therapeutic target to reduce A*β* plaques in the brain and prevent the onset and progression of AD.

Thus far, drug design efforts for BACE1 have been unsuccessful. The earliest BACE1 inhibitors were designed as analogs of existing BACE1 substrates.^5^ Due to the size of the BACE1 active site, these inhibitors had too large a molecular size (*>* 500 KDa) to permeate through the blood-brain barrier (BBB).^5,6^ Therefore, even if they bound to BACE1 with a high affinity *in vitro*, they failed to inhibit the protein *in vivo* within the human brain.

More recent drug design approaches for BACE1 have attempted to resolve the issue of low BBB permeability through small molecule inhibitors. High throughput screening (HTS) tests extensive libraries of small molecules against a target protein—typically through computational methods like molecular docking simulations—to identify hit compounds with desirable bioactivity.^7,8^ Molecular scaffolding leverages a core molecular structure with desirable bioactive properties as a blueprint for designing an inhibitor.^9–12^ Fragment-based drug design (FBDD) combines molecular fragments—tiny molecules identified through HTS—into a full-sized inhibitor.^7,10,13^ These methods have yielded five small molecule candidate drugs -Verubecestat, Lanabecestat, Atabecestat, Elenbecestat, and CNP520—that made it to Phase II/III clinical trials. However, all of these trials were ultimately terminated due to a lack of cognitive improvement in patients and adverse effects in the liver.^14^ Hence, current drug design methods have failed to produce an effective BACE1 inhibitor as an FDA-approved drug for AD.^3^

The challenges in developing an effective BACE1 inhibitor arise from the high dimensionality of the chemical space. The total size of the synthetically accessible chemical space is estimated to be between 10^33^ and 10^60^ molecules, and only a small fraction of these molecules have been explored so far.^15–17^ This sheer magnitude of the chemical space has translated into two key limitations in current methods for designing drugs and BACE1 inhibitors. First, these methods, which include analog development, HTS, molecular scaffolding, and FBDD, require significant amounts of trial-and-error, making them time- and cost-expensive. ^18^ On average, an estimated 10-15 years and $2.8 billion are necessary to discover an effective chemical inhibitor and bring it to the market as a drug.^19,20^ Second, these methods are heavily guided by human knowledge and intuition.^18^ They cannot explore beyond the small region of the chemical space confined within human knowledge, and they can only discover molecules with similar chemical structures and properties to existing molecules. As a result of these issues, an estimated 98% of drug candidates fail before the final stage of the drug development process.^21^

The latest breakthroughs in artificial intelligence (AI) have enabled new approaches to *de novo* drug discovery that resolve the issues of limited human knowledge and trial-and-error. Generative AI leverages deep learning algorithms to understand the complex patterns behind datasets that are incomprehensible to humans. With these patterns, they recreate the statistical distributions underlying these datasets and sample random points from the distributions to generate new samples with similar properties to the initial training dataset.^22^ When applied to molecules, generative AI can generate millions of new compounds and inhibitors with similar bioactive and pharmacological properties to existing inhibitors. Thus, generative AI can efficiently explore a targeted region of the chemical space, drastically outperforming human-driven methods.^18,23^ Through various deep learning architectures, such as recurrent neural networks (RNNs), variational autoencoders (VAEs), and generative adversarial networks (GANs), generative AI has already generated compounds to inhibit a variety of target proteins. ^24–33^ Many of these compounds have displayed promising efficacy during subsequent steps of the drug design pipeline, like *in silico* and *in vitro* testing, demonstrating the effectiveness of this new drug discovery paradigm.^18^

This work applies generative AI to the *de novo* discovery of BACE1 inhibitors. In a novel generative AI framework, the SELFIES molecular notation, an autoencoder model, and a WGAN-GP generator model are implemented to generate inhibitors with similar molecular properties to existing BACE1 inhibitors, exploring the currently understood chemical space much more efficiently than human-driven methods. A Genetic Algorithm is paired with the generator model to optimize the inhibitory activity and BBB permeability of generated inhibitors, exploring previously inaccessible regions of the chemical space to improve upon the predominant shortcomings of current BACE1 inhibitors.

Generative artificial intelligence (Gen-AI) is a relatively new tool for drug discovery and development. However, training Gen-AI models require availability of large data sets. BACE1 has been targeted for the development of drugs for the treatment of neurodegenerative disorders, particularly Alzheimer’s disease but with very limited clinical success. The extensive studies carried out by several research groups over the years generated large quantities of data on to structure activity relationships (SAR) of BACE1 inhibitors. We therefore used BACE1 inhibition as a test case for the use of Gen-AI in structure activity and structure properties relationship studies to discover hitherto unknown BACE1 inhibitors. In short, compound scoring and molecular docking methods confirm that our framework successfully discovers novel candidate drugs for BACE1 and AD.

## Methods

### 1. Dataset

Two molecular datasets are constructed from the ChEMBL 33 database. ^34^ The first dataset, a general dataset representing the explored chemical space, contains all molecular entries in the ChEMBL 33 database (2,372,674 total molecules) and their Simplified Molecular Input Line Entry System (SMILES) notations. SMILES representations leverage one dimensional strings and a specific set of syntactic rules to capture the structural information of molecules.^35^ Due to their simplicity, SMILES strings have become the most popular line notation for representing molecules. ^36^ The second dataset, a targeted dataset, contains all reported BACE1 inhibitors in the ChEMBL 33 database (10,619 total molecules), along with their SMILES notations and IC50 values toward BACE1. By measuring the concentration of a compound required to inhibit the biological function of a target protein by half, IC50 capture potency of an inhibitor toward a target protein.

### 2. Data Preprocessing

Both datasets are preprocessed according to the following steps.

1. All molecules containing a period (”.”) in their SMILES notations, which represents a disconnection, are removed.
2. Duplicate entries (molecules sharing the same SMILES notation) are removed, and in the case of the targeted dataset, their IC50 values are averaged.
3. For the targeted dataset, the IC50 values are converted to pIC50 values, the negative logarithm of IC50, because pIC50 values are easier to model and analyze. ^37^
4. Additional molecular property values are calculated for each molecule in the datasets. Molecular weight (MW) and lipophilicity (represented by the octanol-water partition coefficient, LogP) are two properties in Lipinski’s Rule of Five, which states that candidate drugs with a MW less than 500 Da and a LogP no greater than 5 have better oral bioavailability and membrane permeability through membranes like the BBB.^38^ The Quantitative Estimate of Drug-Likeness (QED) metric measures the drug-likeness of a compound based on its physicochemical properties. ^39^ QED accounts for various factors that determine how the human body processes a compound, including absorption, distribution, metabolism, excretion, and toxicity. The Synthetic Accessibility Score (SAS) metric evaluates the ease with which a compound can be chemically synthesized.^40^ MW, LogP, QED, and SAS are all important metrics to determine the efficacy of a candidate drug.
5. The SMILES notations of molecules are converted to the Self-Referencing Embedded Strings (SELFIES) molecular representation. A major limitation of the SMILES notation lies in the fact that a substantial fraction of SMILES strings do not correspond to valid molecules. This proves to be a serious issue for *de novo* generation of molecules, as many generated SMILES strings do not represent valid molecules. To resolve this issue, Krenn, et al. designed the SELFIES notation, a 100% robust molecular representation through which all strings correspond to a valid molecule. ^41^ The SELFIES notation has only been applied to a limited number of drug discovery models, none of which incorporate a WGAN-GP generative model. ^28,42–44^ This work utilizes the SELFIES notation for *de novo* drug discovery to generate molecules with robust chemical validity.
6. Molecules with SELFIES strings longer than 100 characters are removed from the datasets. This is because longer strings are more difficult to train on in the deep learning models used in this generative AI framework.

After preprocessing, the general dataset contains 2,259,380 molecules and the targeted dataset contains 7,223 molecules. 100,000 and 500,000 molecules are randomly sampled from the general dataset to construct two subsets, a 100k general dataset and a 500k general dataset. These datasets represent the explored chemical space with a smaller number of datapoints, so they are more manageable training datasets for subsequent deep learning models.

### 3. Autoencoder

While SELFIES strings provide a simple and robust molecular representation, they are constructed from distinct alphabetical characters, making them discrete data representations. On the other hand, generative AI algorithms require training data with continuous representations since they create a continuous distribution to model the data. Thus, an autoencoder model is developed to convert SELFIES strings into continuous vector representations of molecules for subsequent use in the generative AI model.

The autoencoder architecture is adopted from Abbasi, et al.,^45^ who used it to successfully convert SMILES strings into vector representations of molecules. In Figure 1a, the implementation of the autoencoder consists of two separate models, an encoder and a decoder, that train simultaneously. The encoder model simplifies a SELFIES string into a 256 length vector in the latent space. It consists of two bidirectional long short-term memory (LSTM) layers, each with 512 units, and a third dense layer of 256 units that outputs the predicted latent vector. LSTMs are advantageous for sequential data forms like strings because they have internal memory mechanisms to remember previous inputs. Following the encoder, the decoder model converts the latent vectors back into their original SELFIES strings. Similar to the encoder, the decoder uses two bidirectional LSTM layers, each with 512 units, and a third dense layer that individually outputs each character in the predicted SELFIES string. In addition, it implements the teacher forcing method, feeding ground truth samples into the decoder model as a secondary input, to prevent error from propagating throughout the predicted SELFIES strings.

**Figure 1:**
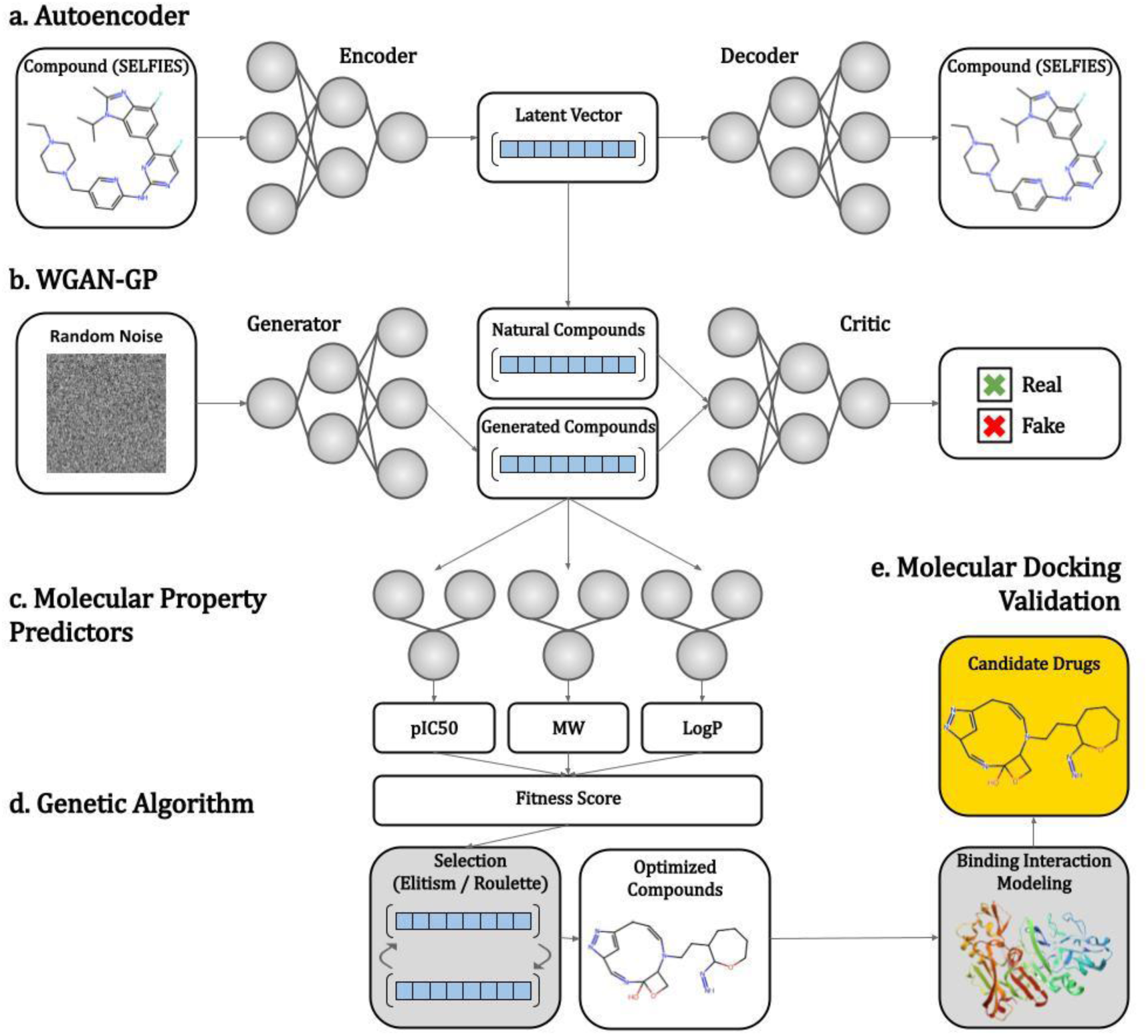
Flowchart of the generative AI framework.

The autoencoder model is trained on the 500k general dataset for 100 epochs at a learning rate of 0.001. During training, the autoencoder learns to perfectly reconstruct the input SELFIES strings of the encoder model into the output SELFIES strings of the decoder model. Through this process, it learns latent vector representations that successfully capture the structural information of molecules.

Using the same model hyperparameters and training dataset, the autoencoder model is trained on the SMILES notation and SELFIES notation to compare between the two string representations of molecules.

### 4. Molecular Property Predictors

In Figure 1c, molecular property predictors are trained to predict three bioactive and pharmacological properties of compounds: pIC50, molecular weight, and LogP.

Since IC50 and pIC50 measure the quantity of a compound required to inhibit the activity of a target protein by half, they are important metrics for determining the potency and efficacy of an inhibitor. However, these values can only be experimentally measured through wet lab techniques, making it impossible to computationally determine the pIC50 of inhibitors.^46^ Thus, a neural network is trained as a quantitative structure-activity relationship (QSAR) model to predict the pIC50 of inhibitors against BACE1.

Traditionally, molecular descriptors have been used to represent molecules in QSAR models.^47^ Molecular descriptors are discrete vector representations that encode a molecule. Their individual bits each signify the absence (0) or presence (1) of a physicochemical, structural, or topological property of a compound. ^36^ Two of the most common molecular descriptors are the Molecular ACCess System (MACCS) keys and the Morgan fingerprints. MACCS keys are 166-bit vectors in which bits denote the presence of structural fragments in a molecule. Morgan fingerprints are 1024-bit vectors in which bits denote the presence of circular substructures around the atoms in a molecule. These two molecular descriptors were compared to the latent vector representations to determine the best molecule input format for QSAR models.

For each molecular representation, a feed-forward neural network is optimized for the number of hidden layers, the number of nodes per layer, and a learning rate through a grid search algorithm. Then, the network is trained on the targeted dataset to predict the pIC50 of compounds toward BACE1. The root-mean-square error (RMSE) values of the three networks are compared to determine the best molecular representation for QSAR models.

Using the best molecular representation and optimal neural network hyperparameters, two additional models are trained to predict the MW and LogP of compounds using the 500k general dataset. Although MW and LogP can be computationally calculated, they are still predicted by QSAR models in this framework to complement the Genetic Algorithm, which operates in the latent space.

### 5. Wasserstein GAN with Gradient Penalty

After molecules in the 100k general dataset and the targeted dataset are converted from SELFIES strings to latent vector representations, a generative AI model is trained to learn the continuous latent space distributions of both datasets. The specific generative model used in this work is the Wasserstein GAN with Gradient Penalty (WGAN-GP). Recently, GANs have grown very popular as generative models for the high quality of their generated samples.^48^ In addition, WGAN-GPs have been shown to be the most robust variant of the GAN, avoiding many issues like mode collapse and non-convergence that are present in standard GANs.^49^ Hence, a WGANGP is selected as the most promising model for this task of *de novo* drug generation.

In Figure 1b, the WGAN-GP consists of two neural networks, a generator model and a critic model, that train simultaneously. The goal of the generator is to convert an input of random Gaussian noise into an output generated compound. The goal of the critic is to take “real” molecules from the training dataset and “fake” (generated) molecules from the generator and to classify them as “real” or “fake”. During training, the two models compete in a zero-sum game, iteratively improving each other through a loss value. The loss value measures the difference between the distribution of “real” molecules and generated molecules through the Earth Mover’s Distance, which is ensured to be finite and well-defined through a gradient penalty. As the loss value updates the weights in both models, the generator outputs more “realistic” molecules to fool the critic while the critic learns to better distinguish between “real” from the training dataset and “fake” molecules that are generated by the generator. Eventually, when both models converge during training, the generator successfully fools the critic with its generated compounds and learns the distribution of the training data.

The generator network contains an input layer of 64 units (to accept a 64-length vector of random Gaussian noise), a hidden dense layer of 128 units, three hidden dense layers of 256 units, and an output dense layer of 256 units that returns a 256 length vector representation of a molecule. The last layer has no activation function, and the other layers contain the Leaky-Relu activation function. In addition, batch normalization, a procedure to normalize data to zero mean and unit variance for improved training, is also executed between the layers. The critic contains three dense layers of 256 units; the first two layers have the Leaky-Relu activation function and the last layer has no activation function.

Once the WGAN-GP is trained to learn the distributions underlying both datasets, it samples random points from these learned distributions to generate new compounds. This is achieved by inputting random Gaussian noise into the generator model and converting the output latent vector into a SELFIES string through the decoder model.

### 6. Optimization with Genetic Algorithm

The WGAN-GP is paired with a Genetic Algorithm (GA) to optimize the bioactive and pharmacological properties of generated inhibitors. Inspired by the process of natural selection, GAs start with a population of candidate solutions and apply the principles of selection, crossover, and mutation to converge toward an optimal solution over an iterative series of evolutions.^50^ In the context of drug discovery, GAs take an initial population of generated inhibitors and perform selection operations to iteratively improve their bioactive and pharmacological properties.^45,51–54^ This enables the GA to explore previously inaccessible regions of the chemical space and generate inhibitors with improved efficacy.

GA optimization is crucial for discovering BACE1 inhibitors since previous inhibitors have encountered many issues with BBB permeability. The GA in this work optimizes the pIC50, MW, and LogP of generated compounds, improving their potency and BBB permeability. In Figure 1d, the GA operates during WGAN-GP training in the latent space, using the selection operation to iteratively improve training dataset molecules and generated compounds. Every n epochs of the WGAN-GP training (*n* = 100 for the 100k general dataset training and *n* = 5 for the targeted dataset training), training molecules and newly generated molecules are screened for their pIC50, MW, and LogP through the molecular property predictors. A fitness score is calculated for each molecule according to Equation 1.

Then, the bottom 10% of molecules in the training dataset by fitness score are replaced by generated molecules through one of two selection schemes: elitism or roulette. In the elitism scheme, the generated molecules with the highest fitness scores are selected for selection. In the roulette scheme, the generated molecules are randomly selected with a probability proportional to their fitness score: the higher the molecule’s fitness, the greater chance it has of being selected. As selection is continually performed, the molecular property distributions of the training dataset are improved. At the same time, the WGAN-GP learns these new distributions and generates new compounds that exhibit improved molecular properties.

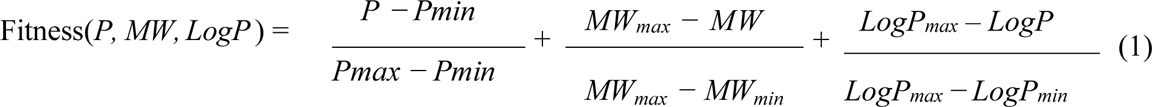

> P represents pIC50. Molecules with higher pIC50, lower MW, and lower LogP values are calculated to have higher fitness values. For a metric *M*, *M_max_* and *M_min_*are the maximum and minimum values of the metric among the training dataset molecules.

Since elitism and roulette are two of the most common selection schemes in GAs, this work implements and compares both of them in the application of molecular optimization. ^55^

### 7. Assessment of Candidate Compounds

After compounds and inhibitors are discovered by the WGAN-GP and GA models, they are assessed for their chemical, pharmacological, and bioactive properties. The SAS and QED values of these compounds are calculated through the python RDKit library, an open source python library for chemoinformatics. ^56^ The pIC50 values of these compounds are predicted by the molecular property predictor model. Since SAS measures synthetic accessibility, QED measures drug-likeness, and pIC50 measures inhibitory activity, compounds with lower SAS values, higher QED values, and higher pIC50 values are easier to synthesize into compounds, better processed by the human body, and more potent inhibitors of BACE1. Thus, these three values are used to calculate the Compound Score metric in Equation 2, which captures the overall efficacy of a candidate compound and inhibitor against BACE1. Calculated Compound Scores are compared between generated compounds to determine the top 100 candidate BACE1 compounds for additional testing.

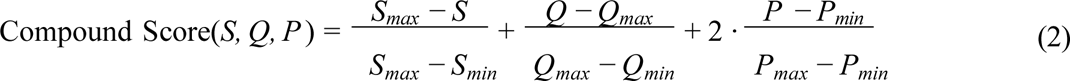

> S represents SAS, Q represents QED, and P represents pIC50. Compounds with lower SAS, higher QED, and higher pIC50 values are calculated to have higher Compound Scores. pIC50 is weighted twice as heavily to prioritze compounds with potent BACE1 inhibition levels. For a metric *M*, *M_max_* and *M_min_* are the maximum and minimum values of the metric among the targeted dataset molecules.

### 8. Molecular Docking Validation

In order to validate the efficacy of the generative AI framework, the 100 generated candidate compounds are compared to Verubecestat, Lanabecestat, Atabecestat, and Elenbecestat, four of the clinical trial-terminated BACE1 drugs. CNP520 is excluded because its molecular information is not present in the ChEMBL 33 database.

In Figure 1e, the validation is conducted through the AutoDock Vina molecular docking simulation, which models the three-dimensional (3D) binding interactions between compounds and BACE1. For the simulation setup, compounds are converted from SELFIES strings into 3D molecular structures containing atomic coordinates. The crystal structure of BACE1 at pH 4.5 (entry 2ZHT in the Protein Data Bank), the most active conformation of the enzyme, is selected as the protein model.^57^ Given these inputs, the simulation utilizes a search algorithm to test various binding conformations of the molecule to the BACE1 active site.^58^ Previous studies have revealed that the BACE1 active site contains two aspartic acid residues, Asp32 and Asp228, that play crucial roles in substrate binding.^59^ Hence, the coordinates of these residues are used to define the simulation search space, and contacts with these residues are used to confirm a valid docking conformation. Residue contacts are determined using the UCSF Chimera molecular visualization software.^60^

The search algorithm returns the optimal binding conformation of each compound that yields the lowest binding energy. Since lower binding energies correspond to more stable binding interactions, molecules with lower binding energies bind more favorably to BACE1. In turn, they act as stronger inhibitors that prevent BACE1 from binding to APP and producing A*β*. Hence, between the generated candidate compounds and the terminated BACE1 drugs, the molecules with the lowest binding energies are determined to have stronger binding interactions with and more potent inhibition of BACE1.

## Results

### 1. Autoencoder

The autoencoder model is trained on both the SMILES and SELFIES representations of molecules in the 500k general dataset to determine the advantages of the SELFIES notation. In theory, the SELFIES notation achieves a more robust molecular representation than the SMILES notation because all SELFIES strings correspond to chemically valid molecules. This robustness comes at the tradeoff of complexity, as SELFIES strings have more complicated vocabularies and grammatical rules compared to SMILES strings. This is evident in Table 1, as SELFIES strings have a similar average length to SMILES strings while using almost 4 times as many unique characters. These unique characters introduce a greater number of underlying patterns for the autoencoder model to learn. Hence, the biggest disadvantage of the SELFIES notation is that it is more difficult for the autoencoder model to train on the SELFIES notation than the SMILES notation.

**Table 1:** Comparison between the syntactic complexity and autoencoder results of SMILES strings versus SELFIES strings.

|  | SMILES | SELFIES |
| --- | --- | --- |
| Average Length (Characters) | 51 | 46 |
| # Unique Characters | 51 | 190 |
| % Perfect Reconstruction | 99.0% | 98.1% |
| % Validity | 99.8% | 100% |

However, the autoencoder model still learns to successfully convert between SELFIES strings and latent vectors. As seen in Table 1, the trained model perfectly reconstructs 98.1% of SELFIES strings in the testing dataset, demonstrating similar accuracy to the SMILES-trained autoencoder. Despite the added complexity of SELFIES strings compared to SMILES strings, SELFIES strings can be represented just as accurately and nearly perfectly by latent vectors. Thus, latent vectors from the trained SELFIES autoencoder become simple and accurate representations of molecules that are always chemically valid. Due to the simplicity and robustness of the latent vectors, they are used to represent molecules in the subsequent models of this framework: the WGAN-GP, the molecular property predictors, and the GA. These models operate in the latent space, inputting and outputting latent vectors, before reconstructing the vectors into SELFIES strings to obtain the final generated and optimized molecules.

### 2. Molecular Property Predictors

Three molecular property predictors are trained on the targeted dataset to predict the pIC50 of compounds toward BACE1. The models use either latent vectors, MACCS Keys, or Morgan Fingerprints to represent molecules. In Table 2, a grid search is performed on each model to optimize the number of hidden layers, units per layer, and learning rate. With these hyperparameters, the models are trained to learn the relationship between a compound’s structure and its inhibitory activity (pIC50) against BACE1. As shown in Table 2, the pIC50 predictor trained on latent vectors achieves the lowest RMSE of 0.58 on the testing dataset, outperforming the other two molecular representations. These results demonstrate that latent vectors provide a better alternative to MACCS Keys and Morgan Fingerprints, the standard molecular representations for QSAR models.

**Table 2:** Comparison between different molecular representations in models to predict the pIC50 of molecules toward BACE1.

| Representation | Layers | Units | Learning Rate | RMSE |
| --- | --- | --- | --- | --- |
| Latent Vector | 1 | 256 | $10^{-4}$ | 0.57 |
| MACCS Keys | 3 | 256 | $10^{-2}$ | 1.46 |
| Morgan Fingerprint | 1 | 64 | $10^{-2}$ | 1.55 |

Latent vector models using the same optimized hyperparameters as found in Table 2 are also trained on the 500k general dataset to predict the MW and LogP of compounds. In Table 3, the MW predictor achieves an RMSE value of 19.93, which equates to only 5% error, and the LogP model achieves an RMSE value of 0.68. In subsequent steps of the generative AI framework, the trained molecular property predictor models are used to evaluate the inhibitory activity, molecular size, and lipophilicity of compounds in the latent space with low error.

**Table 3:** Results of the trained molecular property predictors for MW and LogP.

| Model | RMSE |
| --- | --- |
| MW Predictor | 19.93 |
| LogP Predictor | 0.62 |

### 3. Wasserstein GAN with Gradient Penalty

The WGAN-GP is trained on both the 100k general dataset and the targeted dataset to learn the general molecular property distributions of molecules in the explored chemical space and the specific molecular property distributions of current BACE1 inhibitors. After 10,000 training epochs, the discriminator can no longer distinguish between real samples and generated samples, signifying that the model has converged and the generator has successfully learned the training distributions. Then, by sampling random points from these learned distributions as SELFIES strings, the WGAN-GP generates 250,000 compounds from the general dataset and 250,000 compounds from the targeted dataset.

The WGAN-GP is then assessed based on the validity, uniqueness, and novelty of these generated compounds. Validity is calculated as the percentage of generated SELFIES strings that correspond to chemically valid molecules. Uniqueness is calculated as the percentage of generated SELFIES strings that are not duplicates of each other. Novelty is calculated as the percentage of generated SELFIES strings that are not present in the explored chemical space, as represented by the general dataset. In Table 4, almost all of the generated compounds are valid, unique, and novel. In contrast, previous generative AI models trained on the SMILES notation and similar GAN architectures only achieved validity scores of 30.2% and uniqueness scores lower than 66%.^32,45^ Hence, when the WGAN-GP is trained on the SELFIES notation, it generates a greater number of valid, unique, and novel compounds, discovering new compounds with improved robustness and efficiency.

**Table 4:** The generation statistics of 250,000 generated molecules from the WGAN-GP trained on the 100k general dataset and the targeted dataset.

| Dataset | Validity (%) | Uniqueness (%) | Novelty (%) |
| --- | --- | --- | --- |
| General (100k) | 99.9% | 99.3% | 99.1% |
| Targeted | 100% | 97.5% | 97.3% |

Many of these generated compounds exhibit comparable bioactive and pharmacological properties to the explored chemical space and existing BACE1 inhibitors. Figure 2 illustrates the learned distributions of the WGAN-GP for both datasets. For the general dataset, the WGAN-GP successfully learns and recreates the pIC50, MW, and LogP distributions of the training dataset molecules that represent the explored chemical space. For the targeted dataset, the WGAN-GP recreates the general shapes of these distributions without capturing the finer details at the ends of the distributions. This training limitation is an expected result from the size of the targeted dataset. Since the targeted dataset has over 10-times fewer data points compared to the 100k general dataset, it is more difficult for the WGAN-GP to generalize the molecular patterns behind the molecules in the targeted dataset and to fully explore that region of the chemical space. Overall, however, the WGAN-GP learns to generate inhibitors with similar pIC50, MW, and LogP values to existing molecules and BACE1 inhibitors, discovering candidate BACE1 inhibitors with promising BACE1 inhibition levels and BBB permeability.

**Figure 2:**
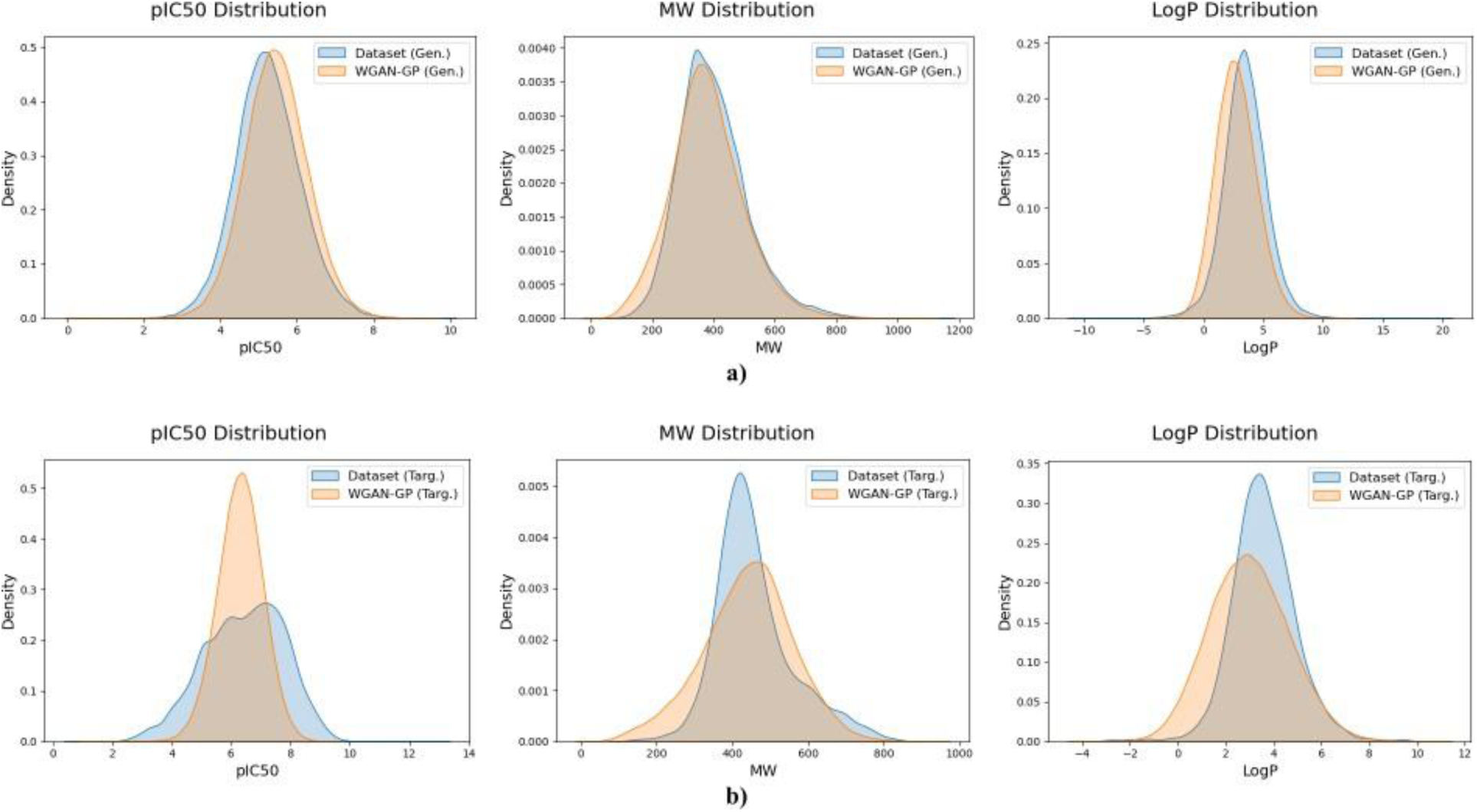
The pIC50, MW, and LogP distributions of 250,000 generated molecules from the WGAN-GP trained on a) the 100k general dataset and b) the targeted dataset.

Lastly, in Table 5, the generated inhibitors from both training datasets are compared for the average values of their pIC50, MW, and LogP values. The general dataset yields compounds with lower MW and LogP values while the targeted dataset yields compounds with higher pIC50 values. Hence, the general dataset is advantageous for generating small molecules with favorable BBB permeability while the targeted dataset is advantageous for generating potent inhibitors of BACE1. This result confirms the effective exploration of the chemical space by the WGAN-GP since current BACE1 inhibitors are known to have reduced BBB permeability but improved BACE1 inhibition compared to other molecules.

**Table 5:** Comparison between the average pIC50, MW, and LogP values of the 250,000 generated molecules from the 100k general dataset and the targeted dataset.

| Property | Dataset | Average |
| --- | --- | --- |
| pIC50 | General (100k) | 5.45 |
|  | Targeted | 6.31 |
| MW | General (100k) | 379.0 |
|  | Targeted | 440.4 |
| LogP | General (100k) | 2.74 |
|  | Targeted | 2.97 |

### 4. Optimization with Genetic Algorithm

In order to improve the inhibitory activity and BBB permeability of generated compounds, a GA is implemented in tandem with the WGAN-GP. Using pIC50, MW, and LogP values, the GA calculates a fitness value for each molecule (according to Equation 1) and iteratively replaces low-fitness training dataset molecules with high-fitness generated molecules. This is achieved through one of two selection schemes, elitism or roulette, which are both implemented in the GA and compared.

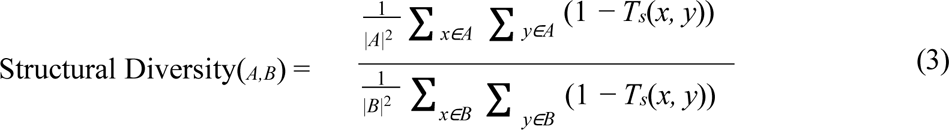

> Relative structural diversity between optimized compounds (set A) and standard generated compounds from the same dataset (set B). A higher structural diversity score (closer to 1) is more desirable.

For both selection schemes, the GA is trained for 5000 epochs until the average fitness of the molecules in the training dataset stagnates. Then, 10,000 compounds are generated from each selection scheme. In Table 6, these compounds are compared based on structural diversity scores, novelty scores, and the percentile of top current BACE1 inhibitors they fall within based on fitness. Structural diversity is calculated according to Equation 3, in which the average structural difference between optimized compounds is divided by the average structural difference between standard generated compounds from the same dataset. The Tanimoto Similarity (TS) metric is leveraged to compute the structural difference between molecules based on their MACCS key representations. ^61^ The structural diversity score ensures that the GA does not over-optimize the fitness of its molecules by converging on a tiny region of the chemical space in which molecules are very similar.

**Table 6:** Results of a sample of 10,000 generated compounds from both selection schemes and for both datasets. Based on their fitness score, the generated compounds are sorted into the top 1% and 5% of current BACE1 inhibitors.

| Dataset / Scheme | Structural Diversity | Novelty (%) | Top 1% | Top 5% |
| --- | --- | --- | --- | --- |
| General (100k) / Elitism | 0.96 | 99.5% | 4,423 | 7,117 |
| General (100k) / Roulette | 0.94 | 95.1% | 1,790 | 4,776 |
| Targeted / Elitism | 0.80 | 98.0% | 2,642 | 3,722 |
| Targeted / Roulette | 0.92 | 99.3% | 2,129 | 3,482 |

For the 100k general dataset, elitism selection generates a greater number of structurally diverse and novel compounds that display comparable fitness scores to the top BACE1 inhibitors. Since the general dataset already covers an extensive region of the chemical space, the targeted approach of elitism selection effectively converges on the optimal region of molecules. However, for the targeted dataset, roulette selection outperforms elitism selection. It generates novel compounds with much higher structural diversity while only slightly compromising their fitness scores. Since the targeted dataset covers a limited region of the chemical space, the randomized element of roulette selection effectively explores this chemical space without over-optimizing and converging on a small subset of highly similar molecules. Hence, this work demonstrates that elitism selection is advantageous for larger datasets while roulette selection is advantageous for smaller datasets.

As the best selection schemes for the general dataset and targeted dataset respectively, the elitism selection GA and roulette selection GA are used to generate 250,000 optimized compounds per dataset. In Figure 3, the pIC50, MW, and LogP distributions of these optimized compounds represent significant improvements from the distributions of the standard generated compounds from the WGAN-GP. By improving the fitness of the training dataset, the GA discovers an unexplored region of the chemical space containing inhibitors with improved inhibitory activity and BBB permeability. Hence, the GA can successfully generate candidate inhibitors that improve upon the primary shortcomings of current BACE1 inhibitors.

**Figure 3:**
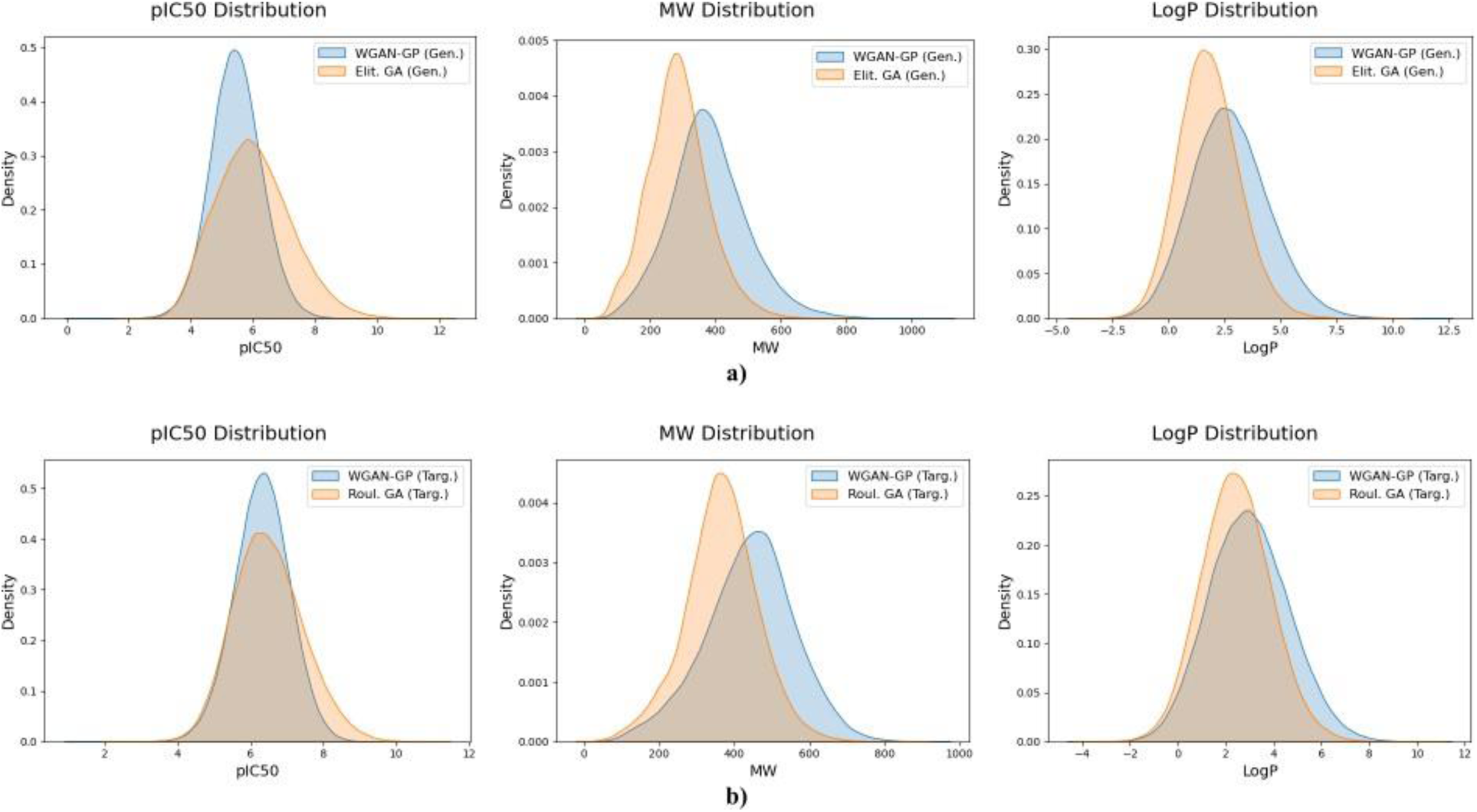
Comparison between the pIC50, MW, and LogP distributions of 250,000 WGAN-GP generated molecules and 250,000 GA generated molecules for a) the 100k general dataset and b) the targeted dataset.

In Table 7, almost all generated compounds from the GA are valid, unique, and novel. Once again, these generation statistics represent an improvement from previous generative AI models that leveraged the SMILES notation.^32,45^ This further proves that the SELFIES notation increases the robustness and efficiency of the generative AI algorithm. In addition, these results demonstrate that the generative AI framework can generate large quantities, on the scale of hundreds of thousands to millions, of novel compounds with comparable bioactive and pharmacological properties to current BACE1 inhibitors. Hence, the WGAN-GP and GA can successfully discover compounds that are candidate inhibitors of BACE1, expediting the drug discovery process.

**Table 7:** The generation statistics of 250,000 optimized molecules from the GA on the 100k general dataset and the targeted dataset.

| Dataset | Validity (%) | Uniqueness (%) | Novelty (%) |
| --- | --- | --- | --- |
| General (100k) | 99.9% | 98.7% | 98.6% |
| Targeted | 99.9% | 98.1% | 98.0% |

### 5. Assessment of Candidate Compounds

Over 1,000,000 total novel compounds are discovered by the generative AI framework between the standard WGAN-GP and GA models trained on both the 100k general dataset and the targeted dataset. To assess the potential of these compounds to perform as BACE1 drugs, their SAS, QED, and pIC50 values are determined. Figure 4 demonstrates that a subset of the compounds have favorable SAS and QED values, meaning that they are easily processed by the human body and easily synthesized in real life. Figures 2 and 3 demonstrate that the generated compounds display similar or improved pIC50 values, or inhibitory potency, compared to existing BACE1 inhibitors.

**Figure 4:**
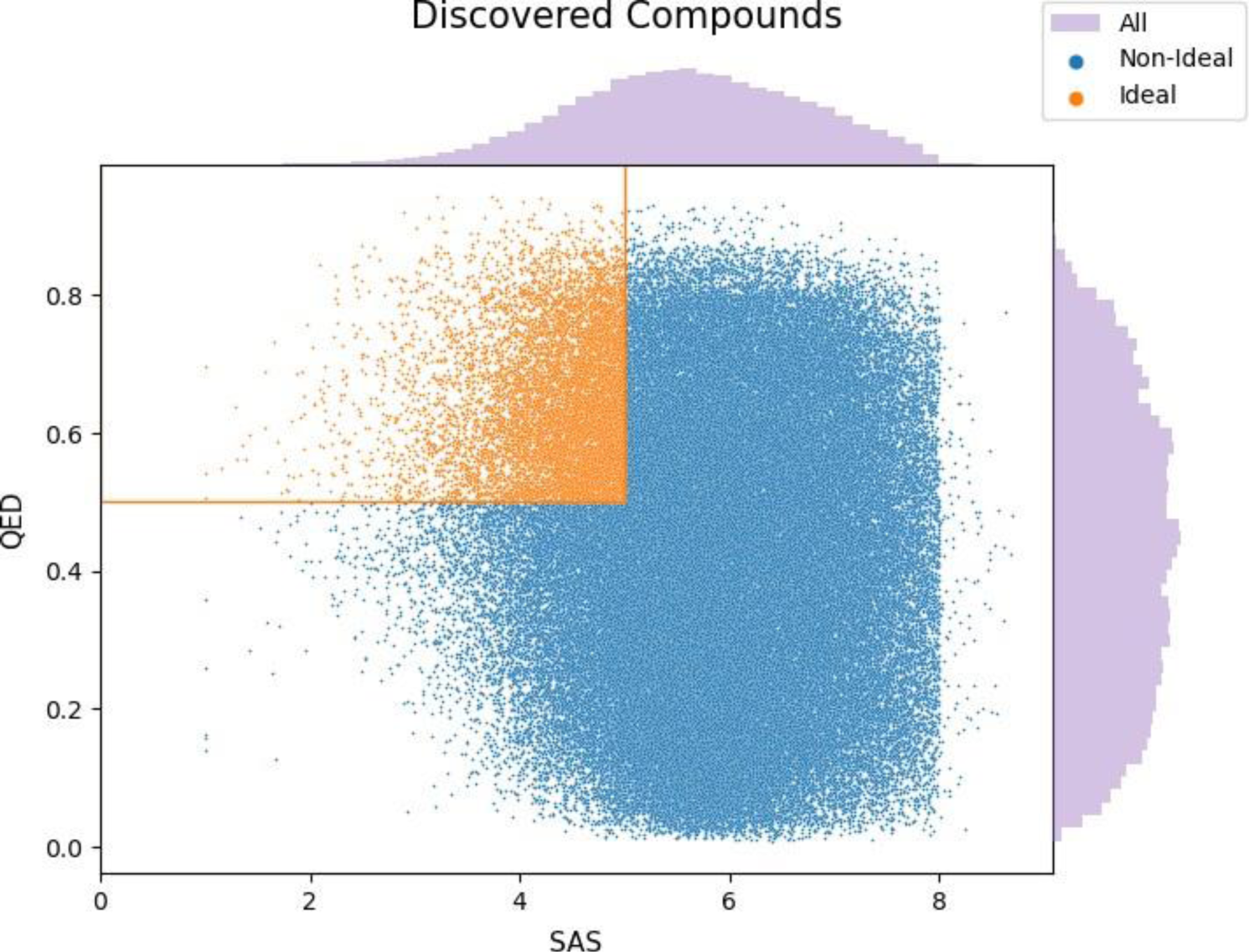
Scatter plot of the QED and SAS values of 10,000 molecules sampled from the optimized compounds from a) the general dataset and b) the targeted dataset. The orange box represents the molecules in the ideal region of drug-like and synthetically accessible chemical space.

Then, using Equation 2, Compound Scores are calculated and used to compare the 1,000,000 generated compounds to the current BACE1 inhibitors. In Table 8, 150 generated compounds perform in the top 1% of existing BACE1 inhibitors based on inhibitory activity, drug-likeness, and synthetic accessibility. This further demonstrates that the *de novo* drug discovery framework successfully generates and optimizes novel compounds that can compete with current BACE1 inhibitors in terms of chemical, pharmacological, and bioactive properties.

**Table 8:**
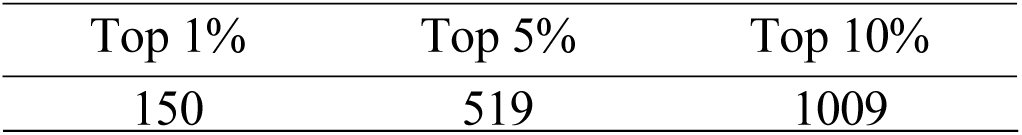
The number of generated molecules with calculated molecule scores in the top 1%, 5%, and 10% of current BACE1 inhibitors.

In addition, Compound Scores are used to rank all 1,000,000 generated compounds and select the top 100 compounds, as shown in Figure 5, as candidate BACE1 compounds for subsequent testing through more robust computational methods.

**Figure 5:**
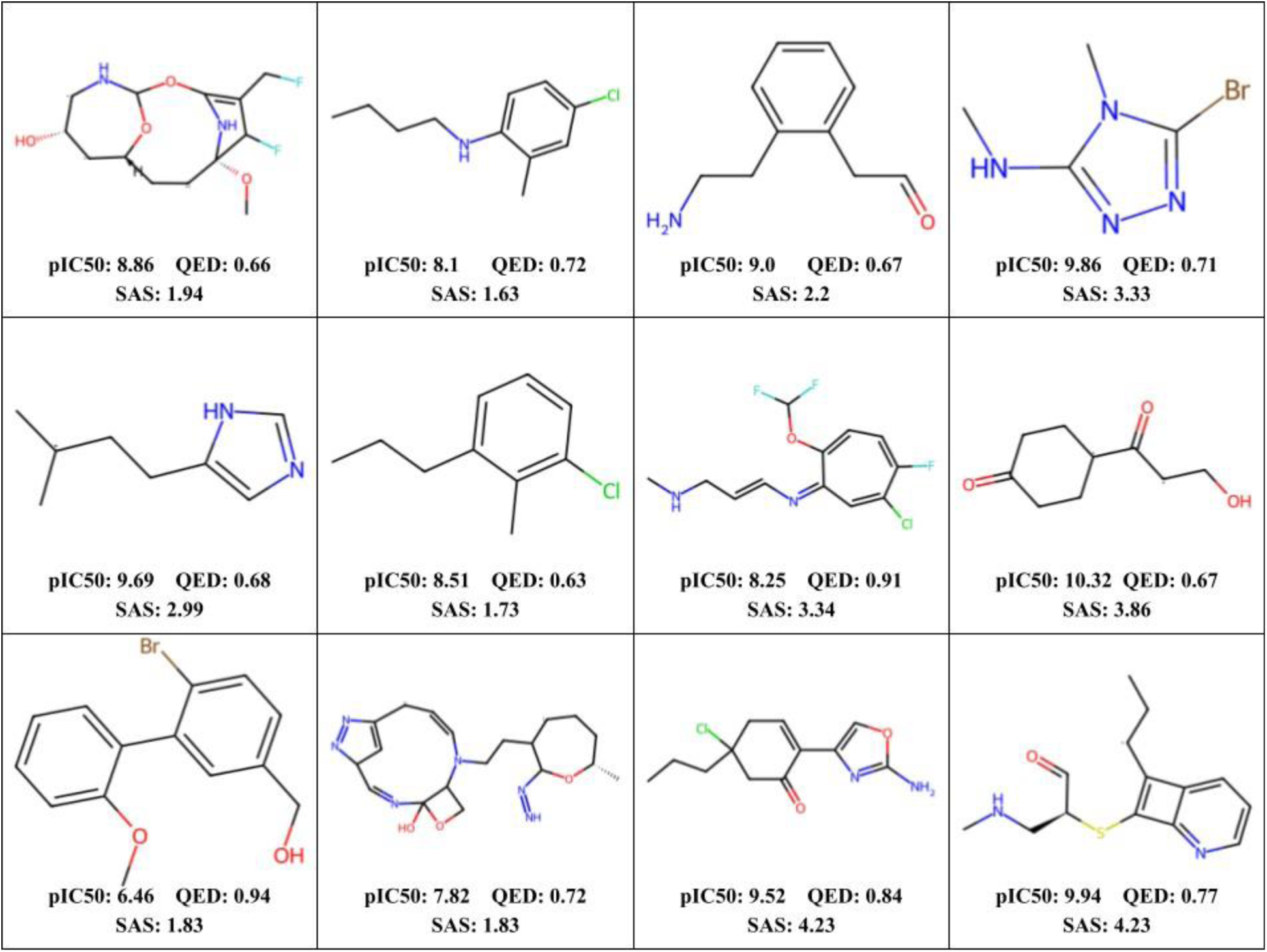
12 randomly selected candidate BACE1 compounds from the top 100 selected compounds, along with their pIC50, QED, and SAS values.

### 6. Molecular Docking Validation

The AutoDock Vina molecular docking simulation is used to compare between the binding interactions of the 100 candidate BACE1 compounds and the binding interactions of the four terminated BACE1 drugs to the BACE1 active site. In Table 9, three of the generated candidate compounds exhibit more favorable binding energies than all four terminated BACE1 drugs. In their specific binding interactions, as shown in Figure 6, they form contacts with Asp32 or Asp228, the key BACE1 binding residues. This signifies a proper interaction with the BACE1 active site to inhibit APP binding and A*β* production. In addition, in Table 9, eleven other generated candidate compounds exhibit more favorable binding energies than at least one of the terminated BACE1 drugs while maintaining proper residue contacts. By forming stronger binding interactions to the BACE1 active site than the terminated BACE1 drugs, these fourteen candidate compounds are promising BACE1 inhibitors and candidate BACE1 drugs. Hence, they can immediately be tested in subsequent steps of the drug design pipeline, verifying the efficacy of the generative AI framework for the *de novo* discovery of BACE1 drugs and for the acceleration of the drug design process.

**Figure 6:**
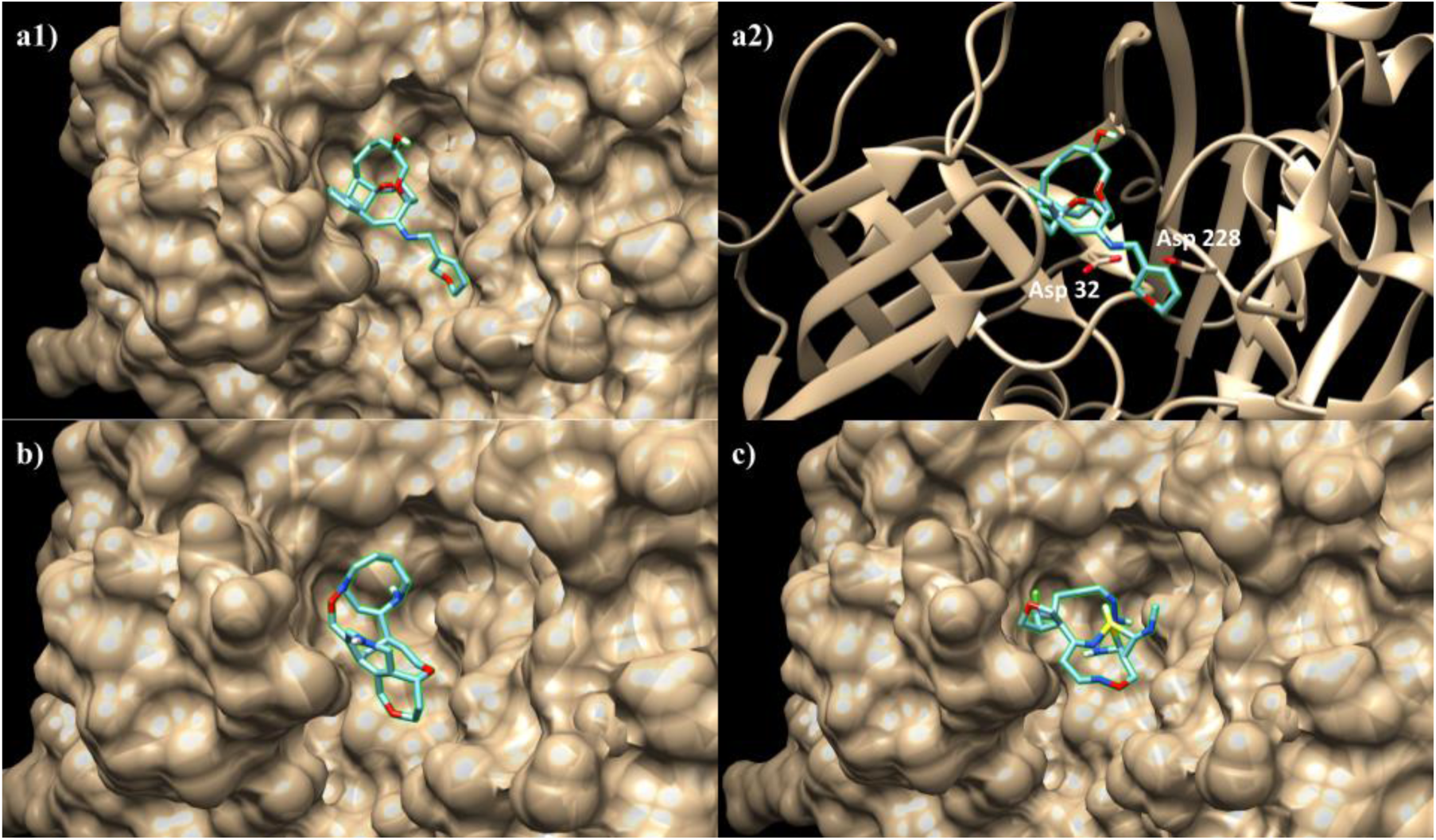
The top simulated poses of **a)** compound 50, **b)** compound 67, and **c)** compound 6 in the BACE1 active site. The Asp32 and Asp228 residues are shown in **a2)** to illustrate the compound’s contacts with BACE1.

**Table 9:** Binding energies and Asp32/Asp228 contacts of top generated compounds, ATABECESTA, ELENBECESTAT, LANABECESTAT, and VERUBECESTAT. Only the compounds with more favorable binding energies than at least one of the candidate BACE1 drugs are shown.

| Compound Name | Binding Energy (kcal/mol) | Contact w/ Asp32 | Contact w/ Asp228 |
| --- | --- | --- | --- |
| Generated Molecule 48 | -9.5 | No | No |
| Generated Molecule 50 | -9.0 | Yes | Yes |
| Generated Molecule 67 | -8.7 | Yes | Yes |
| Generated Molecule 6 | -8.7 | Yes | No |
| LANABECESTAT | -8.3 | — | — |
| Generated Molecule 20 | -7.9 | Yes | No |
| Generated Molecule 90 | -7.9 | Yes | Yes |
| Generated Molecule 53 | -7.9 | Yes | Yes |
| Generated Molecule 43 | -7.8 | Yes | No |
| Generated Molecule 46 | -7.8 | Yes | Yes |
| Generated Molecule 30 | -7.8 | Yes | Yes |
| Generated Molecule 88 | -7.8 | Yes | Yes |
| Generated Molecule 70 | -7.8 | No | No |
| ELENBECESTAT | -7.8 | — | — |
| VERUBECESTAT | -7.8 | — | — |
| Generated Molecule 29 | -7.7 | Yes | Yes |
| Generated Molecule 62 | -7.6 | Yes | No |
| Generated Molecule 34 | -7.4 | Yes | Yes |
| Generated Molecule 82 | -7.3 | No | No |
| Generated Molecule 93 | -7.3 | Yes | No |
| Generated Molecule 25 | -7.1 | No | No |
| ATABECESTA | -7.0 | — | — |

## Conclusion and Discussion

The generative AI framework developed in this work represents the first generative AI approach for the *de novo* discovery of BACE1 inhibitors and AD candidate drugs. The SELFIES molecular representation is leveraged for the first time in the context of a generative AI model to prevent a major shortcoming of previous generative drug discovery models: the discovery of invalid chemical compounds. The autoencoder model converts SELFIES strings into 256-length latent vectors, achieving near-perfect reconstruction accuracy. These latent vectors capture important structural information of molecules in a significantly simplified representation, allowing subsequent models to operate in the simplified latent space. In the molecular property predictors, latent vectors outperform MACCS keys and Morgan fingerprints, two popular molecular descriptors, as the basis for predicting the pIC50 of compounds toward BACE1. In the WGAN-GP, the generator learns to recreate the underlying bioactive and pharmacological distributions of training molecules and efficiently generates over 500,000 compounds that are almost entirely valid, unique, and novel. The GA optimizes the inhibitory activity and BBB permeability of generated compounds, improving upon the primary shortcomings of existing BACE1 inhibitors. This is achieved through a fitness function and selection operation. The elitism selection scheme is found to perform better for larger datasets and the roulette selection scheme is found to perform better for smaller datasets. Lastly, the 1,000,000 discovered compounds from the framework are scored based on their chemical, pharmacological, and bioactive properties, and the top 100 candidate compounds are screened in the AutoDock Vina molecular docking simulation. The simulation models the binding interaction between the compounds and the BACE1 active site and validates 14 candidate drugs to have stronger binding interactions than four of the terminated BACE1 drugs.

Two limitations are uncovered in the generative AI framework that present possible areas of focus for future implementations. First, in Figure 3b, the WGAN-GP does not learn the entire molecular property distributions of the current BACE1 inhibitors in the targeted dataset. This results from the small size of the dataset, which totals only 7,223 molecules. Future works could explore methods of data augmentation, like randomized SMILES and SELFIES molecular representations, to increase the number of data points in smaller datasets and improve WGAN-GP training. Second, in Figure 4, a majority of the 1,000,000 discovered compounds do not have favorable SAS and QED values, so they are not highly drug-like and synthetically accessible. Future works could optimize the SAS and QED values of discovered compounds by incorporating these metrics into the GA fitness function.

The results of this work reveal new possibilities for future efforts aimed at developing a novel BACE1 inhibitor and AD drug. The generative AI framework discovers 14 candidate drugs that display desirable chemical, pharmacological, and bioactive properties and exhibit stronger binding interactions to BACE1 compared to the terminated BACE1 drugs. Future works could test these candidate BACE1 drugs through the later steps of the drug design pipeline, which include molecular dynamics simulations, chemical synthesis, *in vitro* assays, and clinical trials.

Lastly, the generative AI framework developed in this work can be adapted to other target proteins that present themselves as major therapeutic targets. Only minor modifications would be required to retrain the pIC50 predictor on a new targeted dataset containing inhibitors of the target protein. Afterward, the generative AI framework could be applied in a similar fashion to efficiently generate and optimize millions of novel candidate compounds for the target protein.

## Data Availability

The training datasets, python scripts, trained models, and generated candidate drugs associated with this work are publicly available at https://github.com/EvX57/GenAI-Drug-Discovery-for-BACE1. All models are implemented using the Tensorflow package in Python for deep learning.^62^

## Acknowledgements

The authors thank the Simons Foundation and the Simons Summer Research Program for their funding and support. The authors would also thank Stony Brook Research Computing and Cyberinfrastructure and the Institute for Advanced Computational Science at Stony Brook University for access to the high-performance SeaWulf computing system, which was made possible by $1.85M in grants from the National Science Foundation (awards 1531492 and 2215987) and matching funds from the the Empire State Development’s Division of Science, Technology and Innovation (NYSTAR) program (contract C210148).

## TOC Graphic

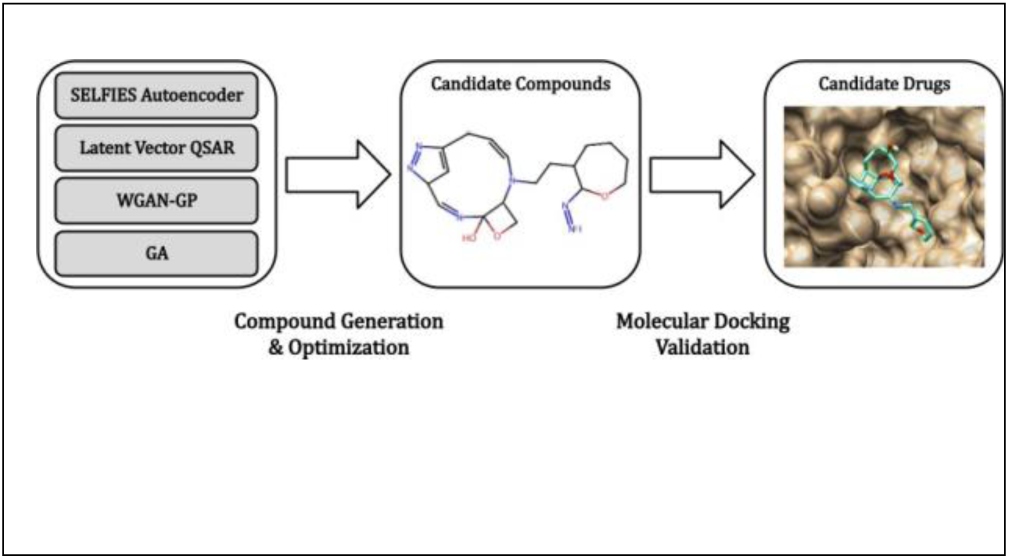

